# Feeding activity of soil oligochaetes under different soil moisture regimes assessed by the bait-lamina test

**DOI:** 10.1101/2025.07.02.662706

**Authors:** Gilda Dell’Ambrogio, Sophie Campiche, Janine W.Y. Wong, Mathieu Renaud, Christina Lüthi, Inge Werner, Benoit J.D. Ferrari

## Abstract

The bait-lamina test is an easy and efficient method to quantify the feeding activity of soil invertebrates, commonly performed under field conditions to measure the impact of chemicals on soils. However, under natural conditions, feeding activity is also influenced by environmental factors such as soil moisture content. The extent of this influence on the results of the bait-lamina test is still poorly described and this can complicate their interpretation. In this study, we optimized the bait-lamina test under laboratory conditions, to assess the influence of soil moisture content on feeding activity of the earthworm *Eisenia andrei* and the enchytraeid *Enchytraeus albidus*, using LUFA 2.2 soil as a substrate. Feeding activity increased linearly with increasing soil moisture until a peak was reached, at a moisture content of 52 % of the maximum water holding capacity (WHC) for earthworms, and 68 % WHC for enchytraeids. Above the optimal moisture content, feeding activity was reduced and was less dependent on soil moisture. The increase in feeding activity up to the optimal moisture content was described by different slopes for the two species. Earthworms consumed the bait faster (on a per unit weight basis) than enchytraeids. Among the two species, the relationship established for *E. albidus* was the more similar to the response obtained in field conditions. Within the range of soil moistures considered by the model, our results demonstrate that feeding activity is positively correlated with soil moisture for two important soil invertebrates, but this increase, as well as the optimal soil moisture content, and the speed of bait consumption are species dependent. The model produced represents a first step for normalizing results from bait lamina tests in the field laying the foundation for future studies to validate its applicability across different soil types and field conditions.

**Highlights:** The optimal soil moisture leading to peak feeding activity was of 52 % and 68 % WHC for *E. andrei* and *E. albidus* respectively.

Earthworms consumed the baits more than three times faster than enchytraeids.

The influence of soil moisture on feeding activity up to the peak can be described by a linear model the slope of which depends on the species tested.

The model obtained for *E. albidus* best represented the field situation.

## 1. Introduction

Soil organisms play an essential role in the breakdown of soil organic matter, which is a fundamental step in the cycling and regulation of nutrients in soil. Among the available tools to assess soil ecological and biological parameters, the standardized bait-lamina method [1] is a simple functional test allowing the *in-situ* measurement of the feeding activity of soil organisms. Feeding activity, an indicator of biological activity, is evaluated at the community level by recording the consumption of organic bait provided in perforated plastic strips inserted vertically into the soil [2,3]. Earthworms and enchytraeids are considered to be the main groups of soil organisms feeding on the bait while the role of micro-arthropods and microorganisms appears to be less important [4–6].

The bait-lamina test can be applied to assess the effect of chemicals on soil organisms, by comparing feeding activity rates between sites of interest and reference sites [7,8], or for the long-term monitoring of soil biological quality [9,10]. The test is nowadays one of the recommended tools for the ecological risk assessment of contaminated soil [11,12]. First developed as a field method, the bait-lamina test has also been described as a promising screening tool for assessing effects of chemicals in the laboratory [7,13–16].

Abiotic factors such as soil moisture and temperature can influence soil faunal activity and therefore complicate the interpretation of the results of the bait-lamina test under field conditions [3,17]. Nevertheless, only a few studies have explored how these factors influence feeding activity. Gongalsky et al. [4] found that feeding activity of enchytraeids increased with increasing soil temperature, while the influence of soil moisture was less evident. Other studies observed that soil moisture was one of the main abiotic factors driving biological activity, suggesting that the first positively influenced the latter [5,18,19]. However, the data generated in these experiments did not allow a detailed characterization of the statistical relationship between the two factors.

In a previous bait-lamina assay performed at the Ecotox Centre [20], a simple increase of feeding activity with increasing soil moisture conditions was observed in an arable field combining different crop types, fertilizer use, and exposure periods. However, the produced model was based on a small number of data points. Additional studies are needed, in order to establish a more robust model, which could be applied to normalize data between tests. Such a model would represent an important tool for comparing bait-lamina test results obtained under field conditions, where soil moisture may vary considerably between sites, and improve the interpretation of field data.

The present study applied the bait-lamina test in the laboratory using two model organisms for ecotoxicity testing, the earthworm *Eisenia andrei* and the enchytraeid *Enchytraeus albidus*. The aim is to investigate the feeding behavior of these two model species, and especially to characterize the relationship between soil moisture content and their feeding activity.

## 2. Material and methods

### 2.1 Preparation of the bait-lamina strips

The bait-lamina PVC strips were 160 mm long, 6 mm wide and 1 mm thick, and perforated with 16 bi-conical apertures 5 mm apart. They were purchased from Terra Protecta (GmbH, Berlin, Germany). The bait substance consisted of a powdery mixture of cellulose, wheat bran flakes and activated charcoal at a ratio of 70:25:5 (w:w:w), which was mixed with Milli-Q water (ratio water to bait of approx. 1.4:1, v:w) to form a paste [1]. Filled bait-lamina strips were air-dried for at least 24h and checked for complete filling of bait-lamina perforations prior to each test.

### 2.2 Test organisms

Based on pilot studies, two species routinely tested in ecotoxicological bioassays were chosen for the bait-lamina tests, i.e., the earthworm (*Eisenia andrei*), and the enchytraeid (*Enchytraeus albidus*). Pilot studies on the collembolan (*Folsomia candida*) were also performed but these showed no bait consumption and the latter organisms was thus not tested further (data not shown).

*Eisenia andrei* Bouché 1972 (Annelida: Oligochaeta) were originally obtained from the farm Lombritonus (Ollon, Switzerland; http://www.lombritonus.ch) and maintained in laboratory cultures. The breeding substrate was composed of a moist mixture of fresh horse manure, composted manure and peat moss in a proportion of 1:1:1 (v:v:v), and fed approximately once a week with finely ground rolled oats previously heated to 105°C for 48h.

*Enchytraeus albidus* Henle 1847 (Annelida: Oligochaeta) were originally obtained from the French National Institute of Agricultural Research (INRA/AgroParisTech, UMR ECOSYS, Versailles, France) and maintained in laboratory cultures. The breeding substrate consisted of a moist mixture of potting soil and LUFA 2.2 at a ratio of 3:2 (v:v). Organisms were fed twice a week with rolled oat flakes and cat food pellets, both previously heated to 105 °C for 48 h and finely ground.

Both cultures were maintained at 20 ± 2°C in the dark for several generations before their use in experiments.

### 2.3 Soil used in experiments

The soil used in all experiments was the standardized LUFA 2.2 soil (LUFA Speyer, Speyer, Germany) a natural sandy loam soil (according to USDA particle size distribution (%)), which had the following properties: pH 5.4, total organic carbon 1.59 %, clay 7.7 %, silt 16.2 %, sand 76.1 % and 45.8 ± 1.9 % maximum water holding capacity (WHC).

In previous pilot studies it was observed that after five days, the bait was partially consumed in LUFA 2.2 soil without the test organisms. To prevent any infestation (community development) by unwanted soil organisms, the soil used for the test lasting 12 days with *E. albidus* was defaunated by freezing at -21 °C for six days prior to test start. For tests lasting only two days (*E. andrei*), the soil was not defaunated because no infestation of the medium was observed in this short lapse of time. To further verify the absence of any additional contribution to bait consumption other than the tested organisms, negative control replicates were run for both species without organisms, as detailed below.

### 2.4 Experimental design

For each experiment, polystyrene plastic containers (171 x 123 x 60 mm) were filled with LUFA 2.2 soil pre-moistened at different range of humidity. Test organisms and bait lamina were then added to the soil.

The soil moisture content of the LUFA 2.2 soil was measured in triplicates by gravimetric method (oven drying at 105°C). Gravimetric moisture content is defined as the amount of water present in the soil, expressed as percentage of soil dry weight. However, the gravimetric moisture content alone does not give sufficient information when it comes to comparison between different soils, because of differences in soil types can lead to differences in water retention. To allow a potential comparison with other soil types, and to facilitate the link with the optimal moisture conditions, usually expressed in standard guidelines as percentage of the maximum WHC, the gravimetric moisture contents measured in these experiments will be expressed as relative moisture contents, i.e., of WHC.

For earthworms, two tests were carried out at the following ranges of nominal relative moisture contents: 20, 40, 60, 80, and 100 % of the WHC and 30, 45, 60, 75, 90, and 105 % of the WHC, using five replicates for each moisture treatment. The nominal 60 % WHC represents standard testing conditions for earthworms in ecotoxicology [21–23] and was thus considered as “reference” to allow comparison with the other moisture treatments. Three negative control replicates were run at two 60% and 100% of the WHC without earthworms, to further verify if the bait was consumed in the absence of the test organisms.

For the enchytraeid experiments, a range of five moisture contents were tested with the following nominal relative moisture contents: 40, 50, 60, 70, and 80 % of the WHC, using four replicates per moisture treatment. The reference moisture content was set at 50 % WHC, representing standard testing conditions for enchytraeids in ecotoxicology [24,25]. Two negative control replicates containing no enchytraeids were included for each moisture treatment.

For each moisture treatment, the gravimetric soil moisture content was measured, as mentioned above, at the beginning and end of the test, based on a composite sample from all replicates.

The number of organisms per replicates was chosen based on the geometric mean of several population densities, collected from the literature for arable soils in Switzerland and neighboring countries (see S1 Table and S2 Table for details). Moreover, to allow a comparison between the feeding rates, a similar biomass, i.e. wet weight of the total number of individuals per replicate, was used for the two species.

For the earthworm experiments, five organisms were placed into 900 g of moist LUFA 2.2 soil for each replicate, corresponding to a field density of 225 individuals m^-2^. Only adults aged between 2 and 8 months and with an individual wet weight between 300 to 600 mg were used for the tests. Based on the range of individual earthworm weight used, the estimated total average biomass of five individuals per replicate was 2.25 ± 0.75 g. The selected worms were acclimated for 48 hours in a separated container filled with LUFA 2.2 soil moistened at 60 % WHC, prior to the tests.

For the enchytraeid experiments, 230 organisms at least 1cm long and of approximately the same size were placed into 600 g of moist soil, corresponding to a field density of 11’000 individuals m^-2^. The wet weight of a single individual was estimated by performing repeated weighing measures of a defined enchytraeid number and corresponded to 0.010 ± 0.003 g. The estimated total average biomass of 230 individuals per replicate was 2.30 ± 0.69 g. *E. albidus* were extracted from the culturing substrate by a wet extraction adapted from the annex A of the ISO guideline 16387 [25]. Briefly, approximately 500 ml of culture soil was carefully crumbled and placed into a metal sieve (180 mm diameter, mesh size 1 to 2 mm). The bottom of the sieve was previously covered with a perforated aluminum foil to avoid soil particles passing through the sieve. The sieve was hung in a plastic bowl, which was filled with tap water until it covered the soil completely. Because of gravity and of their constant motion, the enchytraeids passed through the perforated aluminum foil and the sieve and fell into the water. Water was kept cold with an ice pack and constant aeration was provided with a pump, since a lack of oxygen could cause mortality to the worms [26]. The extraction duration ranged from 6 to 13 h in order to minimize the time the worms spent in the water. Longer periods were found to cause excessive stress in the animals (i.e., lower mobility, higher mortality). At the end of the extraction procedure, the greater part of the water present in the bowl was slowly decanted; the worms were rinsed with tap water and immediately transferred to petri dishes for counting. Before starting the experiment, enchytraeids were acclimated in the test vessels for 48 hours, after which the bait-lamina strips were added.

For both earthworms and enchytraeids experiments, five bait-lamina strips were inserted horizontally into each test container, just beneath the surface of the soil and about 2 cm apart (Fig. 1).

**Fig. 1.**
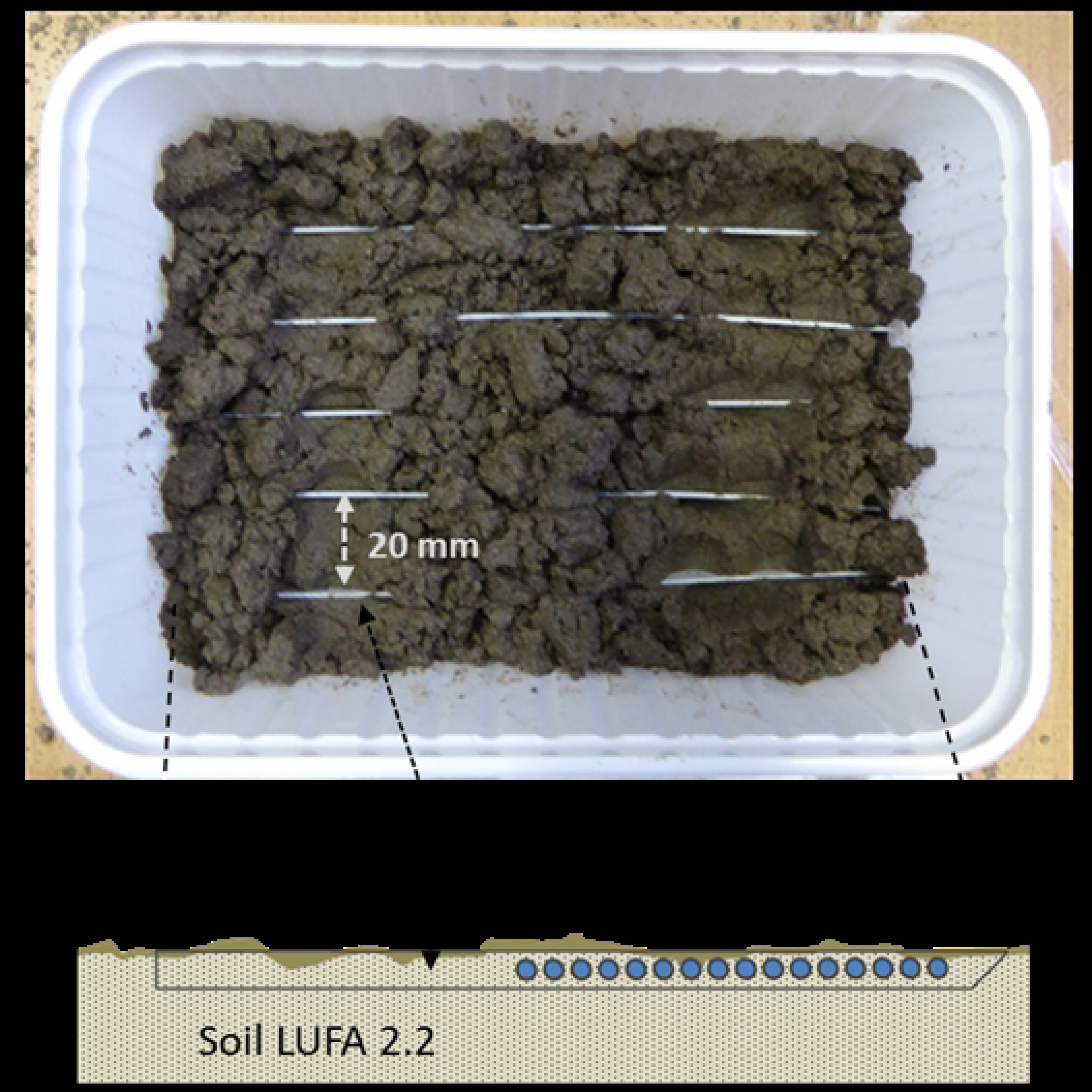
Test system for exposing organisms to five bait-lamina strips.

The test containers were subsequently covered with a 500-µm mesh cotton fabric held in place with a perforated lid in order to prevent escape of the organisms. Test duration was determined in pilot studies and defined as the time required to reach a mean feeding activity of more than 30 % in the reference replicates [1]. This took 2 days for earthworms and 12 days for enchytraeids. All tests were conducted at 20 ± 2 °C with a 16:8-h light-dark photoperiod, according to standard testing and culture conditions for ecotoxicity testing [21–24].

For the enchytraeid test, which had a longer duration compared to earthworms, soil water content was checked by weighing the test containers once a week and Milli-Q water was added when a weight loss of more than 2 % was observed [24]. The moisture content of the earthworm tests was not monitored, as tests only last for 2 days.

At test termination, bait-lamina strips were carefully removed from the soil and gently rinsed with tap water [3,7]. Feeding activity was assessed according to ISO 18311 [1]. Food consumption on the bait-lamina strips was categorized by attributing a score of 2 to a completely empty hole, 1 to a partly empty hole, and 0 to a hole still containing all of the bait. Feeding activity was expressed as percentage of pierced holes per strip (percentage of bait consumed).

Surviving organisms were retrieved from the test soil and their numbers recorded. Living earthworms were extracted by hand, while survival rate of enchytraeids was assessed by wet sieving extraction [27]. Briefly, 3 L of tap water was added to the soil of each replicate, and enchytraeids were fixed with an aqueous solution of 4 % (v:v) formaldehyde. Subsequently, the wet soil was washed 7-8 times through a 355 µm mesh sieve and the extracted enchytraeids were stained with a few mg of Bengal rose and transferred to a petri dish for counting.

### 2.5 Statistical analyses

To allow a better comparison between the different experiments, results of feeding activity for both species were converted into daily feeding activity by dividing the overall feeding activity by the test duration in days. All data were checked for the assumptions of normality and homogeneity of variance by the Shapiro-Wilk and the Levene test, respectively. An analysis of variance followed by a Dunnet’s post hoc test with Holm’s correction for multiple comparisons was used to compare the different treatments with the reference response, and an unpaired two samples t-test was used for a comparison between two treatments only. To investigate the correlation between feeding activity and soil moisture content, Spearman’s test was used. Based on the obtained results, further analyses focused only on data showing a strong and clear correlation, i.e., from lowest to the optimal moisture content, omitting results from treatments above the optimum moisture content of each species. Correlation between daily feeding activity and moisture within this range was analyzed by means of simple and multiple linear regressions. First, two simple linear regressions were fitted, one for each species. Secondly, differences between the two experiments were investigated by means of a multiple linear regression with the fixed effects moisture and species, plus the interaction between moisture and species. Multiple linear regressions were then also used to compare the linear regressions produced in this study with the ones obtained a previous field study from Campiche et al. [20]. From the field study, the maximum WHC of the field soil was not available and moisture contents were given in gravimetric units (percentage of the soil dry weight). Therefore, for this last step, moisture contents were always expressed in gravimetric units for comparison. All statistical tests and linear regression models were performed with the R software within the R studio environment.

## 3. Results

### 3.1. Test performance

Overall mortality rate for the two species was low (< 10 %), except for the earthworm treatment at 25 % WHC, where mortality was 12 % (see SI, Section 2). At the end of the test, average feeding activity in reference moisture treatments met the validity criteria (≥ 30 %) required for the controls in the standard testing guideline [1]. Negative control replicates without test organisms showed no bait consumption at all moisture treatments, except for the nominal 80 % WHC (enchytraeid test) and 100 % WHC (earthworm test), where the average daily feeding activity was 1.04 ± 1.30 and 0.08 ± 0.04 (% of consumed bait per day ± SD), respectively (see S3 Table).

The values of relative soil moisture content measured at the test start were in most cases lower than the nominal values (Table 1). Also, the measured values were lower at the test end compared to the test start (see S4 Fig. and S5 Fig.). Both differences were more pronounced for the earthworm tests compared to the enchytraeid test. Because of these differences, the average values measured between the start and the end of the test were used for the data analysis, as indicated in Table 1. The only exception was the nominal 80 % WHC treatment of the first earthworm test, for which the measured value for relative moisture content was higher at the test end compared to the test start (see Table 1 and S4 Fig. A) and was considered little realistic, since no moisture adjustment was performed for this test. For this specific treatment, the relative moisture content measured at the test start was then considered rather than the average between start and end (see Table 1). For the two tests on earthworms, the results of feeding activity at the two reference moisture contents (nominal 60 % WHC) were not significantly different (unpaired two samples t-test > 0.05). Therefore, the two tests were pooled and the two reference moisture treatments (measured 51 % and 53 % WHC), as well as respective values of feeding activity, were merged as a unique treatment, corresponding to the average of the two, i.e. 52 % WHC.

**Table 1.** Nominal and measured values (at test start and test end) of relative soil moisture content (% of the maximum water holding capacity, WHC, n = 2 for nominal 75 % WHC, Earthworms - test II, and n = 3 for all other replicates ± standard deviation), percentage of difference between nominal and measured (test start), and values used for the data analysis (mean of measured relative moisture contents between the test start and test end. Grey shaded lines indicates the reference moisture conditions considered for the tests. For the negative controls without organisms (Ctrl), the mean was not calculated because not used for the data analysis; n.d. = not determined.

| Nominal relative moisture content (% WHC) | Measured relative moisture content (% WHC ± SD) at test start | Difference in moisture contents between nominal and measured (test start) (%) | Measured relative moisture content (% WHC ± SD) at test end | Relative moisture content used for data analysis (mean between measured start and end, % WHC) |
| --- | --- | --- | --- | --- |
| **Earthworms – test I** |  |  |  |  |
| 20 | 17.76 ± 1.33 | 11 | 14.64 ± 0.19 | 16 |
| 40 | 38.43 ± 0.81 | 4 | 34.61 ± 0.37 | 37 |
| 60 | 58.20 ± 0.57 | 3 | 48.68 ± 6.31 | 52 <sup>a</sup> |
| 80 | 78.65 ± 0.36 | 2 | 87.15 ± 2.18 <sup>b</sup> | 79 |
| 100 | 95.28 ± 0.79 | 5 | 87.33 ± 1.22 | 91 |
| **Earthworms – test II** |  |  |  |  |
| 30 | 25.60 ± 0.46 | 15 | 23.60 ± 0.14 | 25 |
| 45 | 40.50 ± 0.84 | 10 | 38.08 ± 0.22 | **39** |
| 60 | 50.71 ± 0.54 | 15 | 51.80 ± 3.24 | **52<sup>a</sup>** |
| 75 | 71.60 ± 1.03 | 4 | 75.49 ± 0.80 | **74** |
| 90 | 88.39 ± 1.60 | 2 | 92.61 ± 4.68 | **90** |
| 105 | 101.10 ± 2.60 | 4 | 106.51 ± 7.16 | **104** |
| 60 (Ctrl) | 56.31 ± 2.61 | 6 | n.d. | - |
| 100 (Ctrl) | n.d. | n.d. | n.d. | - |
| **Enchytraeids** |  |  |  |  |
| 40 | 40.67 ± 0.36 | 2 | 39.65 ± 0.81 | **40** |
| 40 (Ctrl) | 40.67 ± 0.36 | 2 | 37.26 ± 0.14 | - |
| 50 | 48.83 ± 0.49 | 2 | 49.15 ± 1.11 | **49** |
| 50 (Ctrl) | 48.83 ± 0.49 | 2 | 44.26 ± 0.47 | - |
| 60 | 58.61 ± 0.40 | 2 | 57.64 ± 0.35 | **58** |
| 60 (Ctrl) | 58.61 ± 0.40 | 2 | 58.46 ± 0.22 | - |
| 70 | 69.10 ± 13.23 | 2 | 68.05 ± 1.17 | **68** |
| 70 (Ctrl) | 69.10 ± 13.23 | 2 | 67.16 ± 1.31 | - |
| 80 | 77.45 ± 1.03 | 3 | 77.61 ± 1.77 | **78** |
| 80 (Ctrl) | 77.45 ± 1.03 | 3 | 74.68 ± 0.29 | 4 |
| <sup>a</sup> corresponds to the mean between the moisture contents of the two earthworm tests pooled together (53 and 51).<br><sup>b</sup> was not retained for the mean because unrealistically higher and considered then as outlier. |  |  |  |  |

### 3.2. Soil moisture effect on feeding activity

The results of feeding activity (overall and daily) for the different soil moisture contents (expressed as gravimetric and relative moisture content) are provided in SI2 and depicted in Fig. 2. For the earthworms pooled test (Fig. 2A), the average daily feeding activity was the highest (21.82 ± 10.41 % of consumed bait per day ± SD) at the reference relative moisture content (52 % WHC) and were significantly lower at all other moisture treatments.

**Fig. 2.**
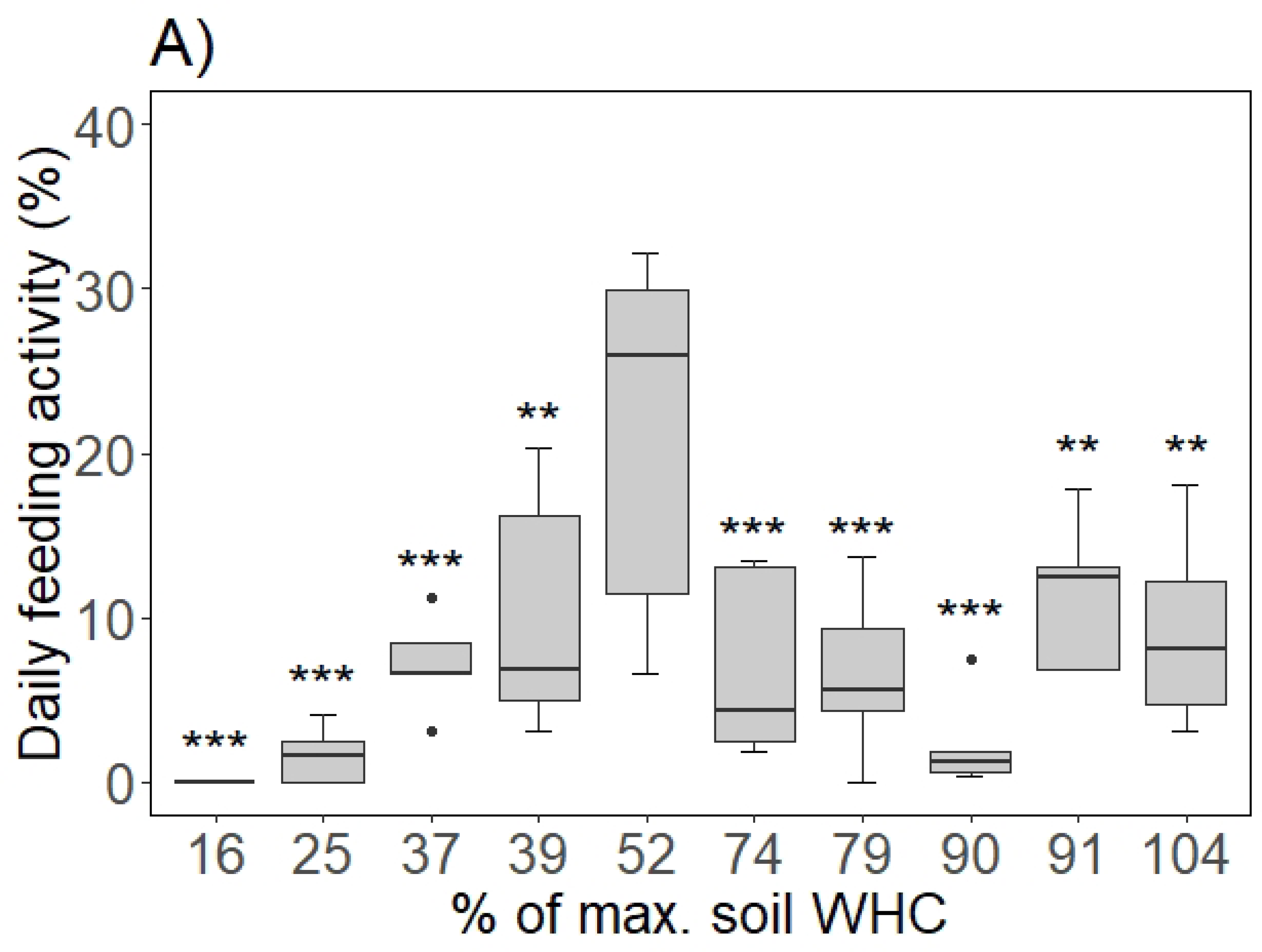
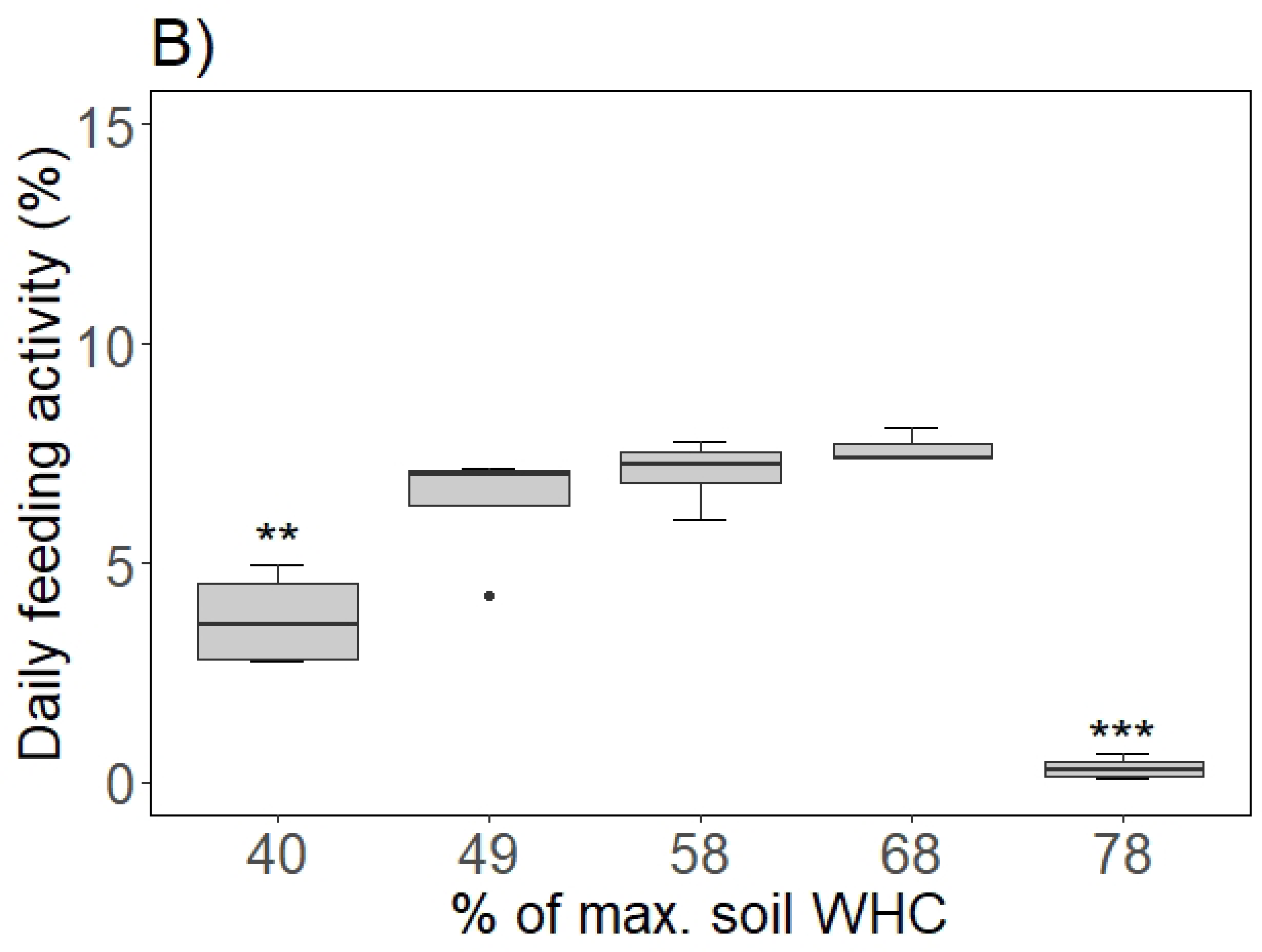
Daily feeding activity (% of consumed bait per day) at different relative soil moisture contents (expressed as % of maximum water holding capacity, WHC) (A) for the earthworm, *E. andrei* (n = 10 at 52 % WHC and n = 5 at all other relative moisture contents), and (B) for the enchytraeid, *E. albidus* (n = 3 at 68 % WHC, n = 4 at all other relative moisture contents). Significant differences from the reference moisture (i.e., 52 % WHC for *E. andrei* and 49 % WHC for *E. albidus*) are indicated by * (p < 0.05), ** (p < 0.01) and *** (p < 0.001), according to Dunnet’s post-hoc with Holm’s correction.

For enchytraeids (Fig. 2B), the highest average daily feeding activity (7.62 ± 0.39 % of consumed bait per day ± SD) occurred at a relative moisture content higher than the reference (68 % rather than 49 % WHC), although the difference in result between the two moisture conditions was not statistically significant. Average daily feeding activity at 40 % WHC and 78 % WHC was significantly lower compared to the reference moisture. At 78 % WHC, most of the enchytraeids were found on the surface of the soil, which was flooded.

The relationship between daily feeding activity and relative soil moisture content until peak feeding activity could be described by a simple linear regression for both test species (Fig. 3). Differently, when considering the whole datasets, no significant correlation was found (Spearman’s tests, p > 0.05). For the relative moisture contents up to the optimum, i.e., from 16 % to 52 % WHC for *E. andrei* and from 40 % to 68 % WHC for *E. albidus*, a multiple linear regression, including two fixed effects (species and moisture) and their interaction, showed a significant influence of the interaction term (p > 0.05), indicating that there are differences in moisture effects on feeding activity between the two species (Table 3).

**Fig. 3.**
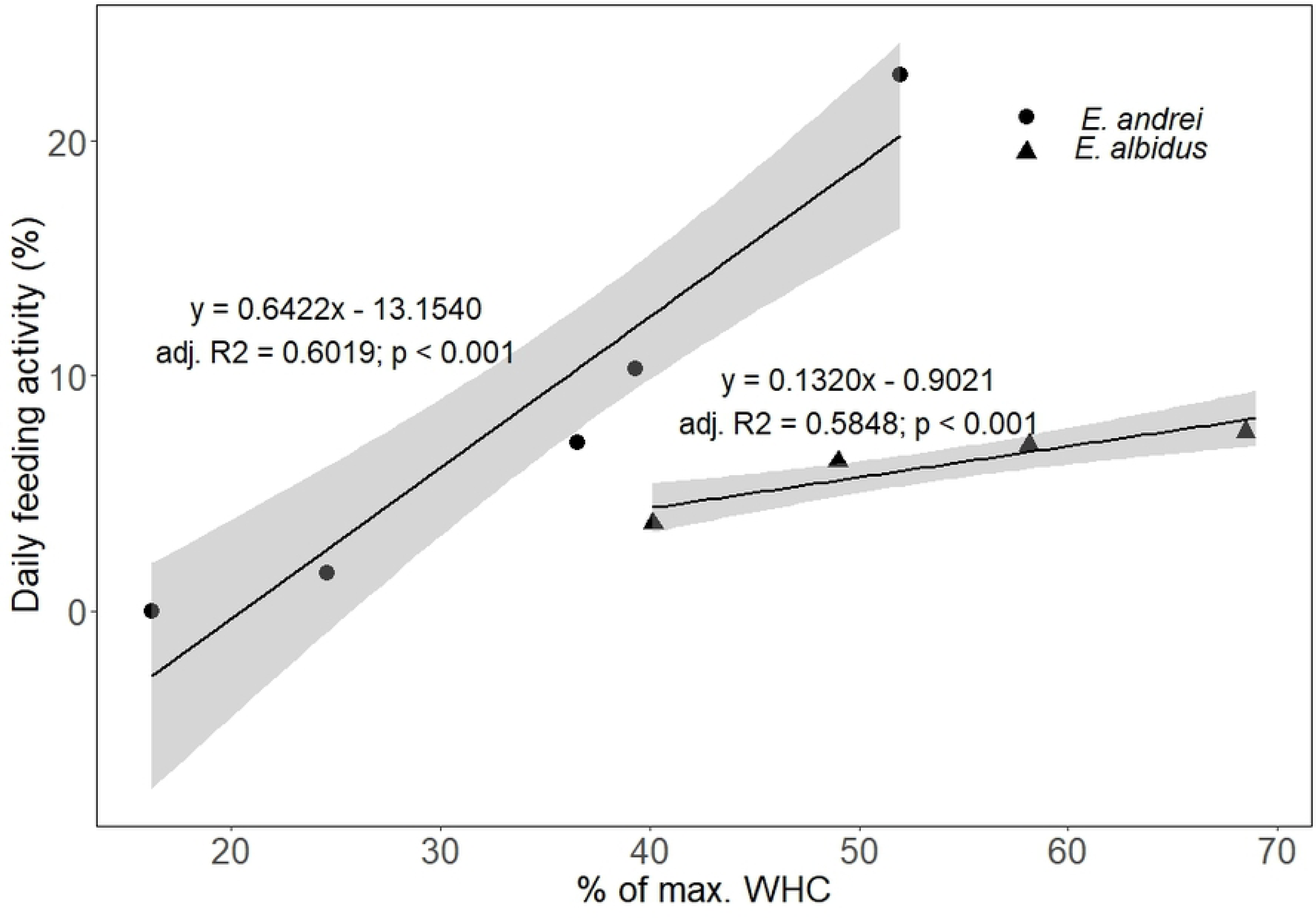
Linear regressions (black lines) and 95% confidence interval (grey shades) of the daily feeding activity (in % of consumed bait per day) of each test species exposed to different soil moistures (% of maximum water holding capacity (WHC)) up to their respective optima. Each symbol (black dots for *Eisenia andrei* and black triangle for *Enchytraeus albidus*) represents the average daily feeding activity per treatment.

**Table 3.**
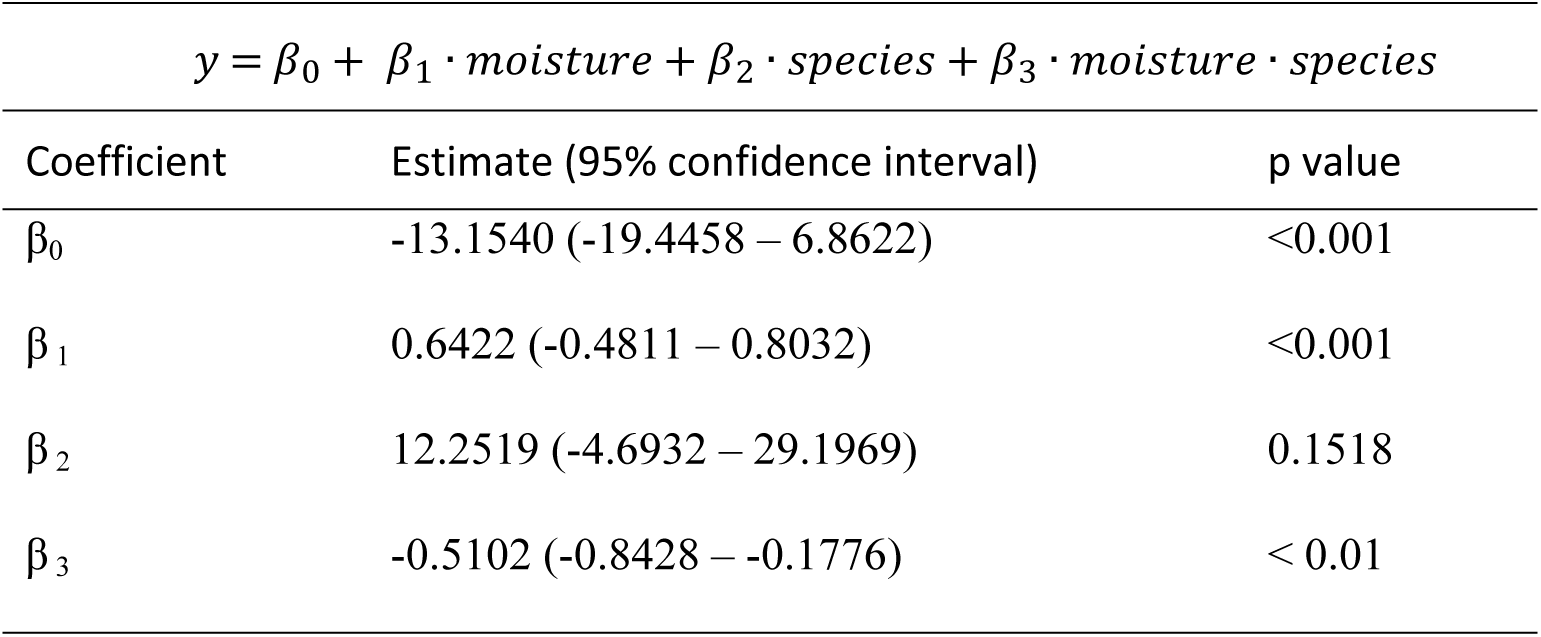
Parameters of the multiple regression analysis for the daily feeding activity response to the two predictors ’moisture’ and ’species’, with interaction between the two factors. Data on moisture are expressed as percentage of the maximum water holding capacity (WHC). The two species compared are *Eisenia andrei* and *Enchytraeus albidus*. β0, β1, β2, and β3 are the coefficients of the linear regression. Adjusted R square = 0.6085.

| $y = \beta_0 + \beta_1 \cdot \text{moisture} + \beta_2 \cdot \text{species} + \beta_3 \cdot \text{moisture} \cdot \text{species}$ | | |
| --- | --- | --- |
| Coefficient | Estimate (95% confidence interval) | p value |
| $\beta_0$ | -13.1540 (-19.4458 – 6.8622) | <0.001 |
| $\beta_1$ | 0.6422 (-0.4811 – 0.8032) | <0.001 |
| $\beta_2$ | 12.2519 (-4.6932 – 29.1969) | 0.1518 |
| $\beta_3$ | -0.5102 (-0.8428 – -0.1776) | < 0.01 |

Finally, multiple regression analysis was performed with the regression model obtained in a previous field experiment (Campiche et al. [20]). The linear regression model produced in the mentioned study is showed in Fig. 4, in addition to the ones obtained in the present studies. For this comparison, moisture content was expressed as gravimetric water content (mass of water per mass of dry soil, in percentage). The results of the multiple regressions (Table 4) showed no significant differences in the feeding activity response to moisture, when comparing *E. albidus* (this study) with Campiche et al. [20], and a common slope of 0.3326 (0.2775 – 0.3879) was estimated for the two studies. On the other hand, a significant difference was found when comparing *E. andrei* (this study) with Campiche et al. [20] with a significant contribution from the interaction term.

**Fig. 4.**
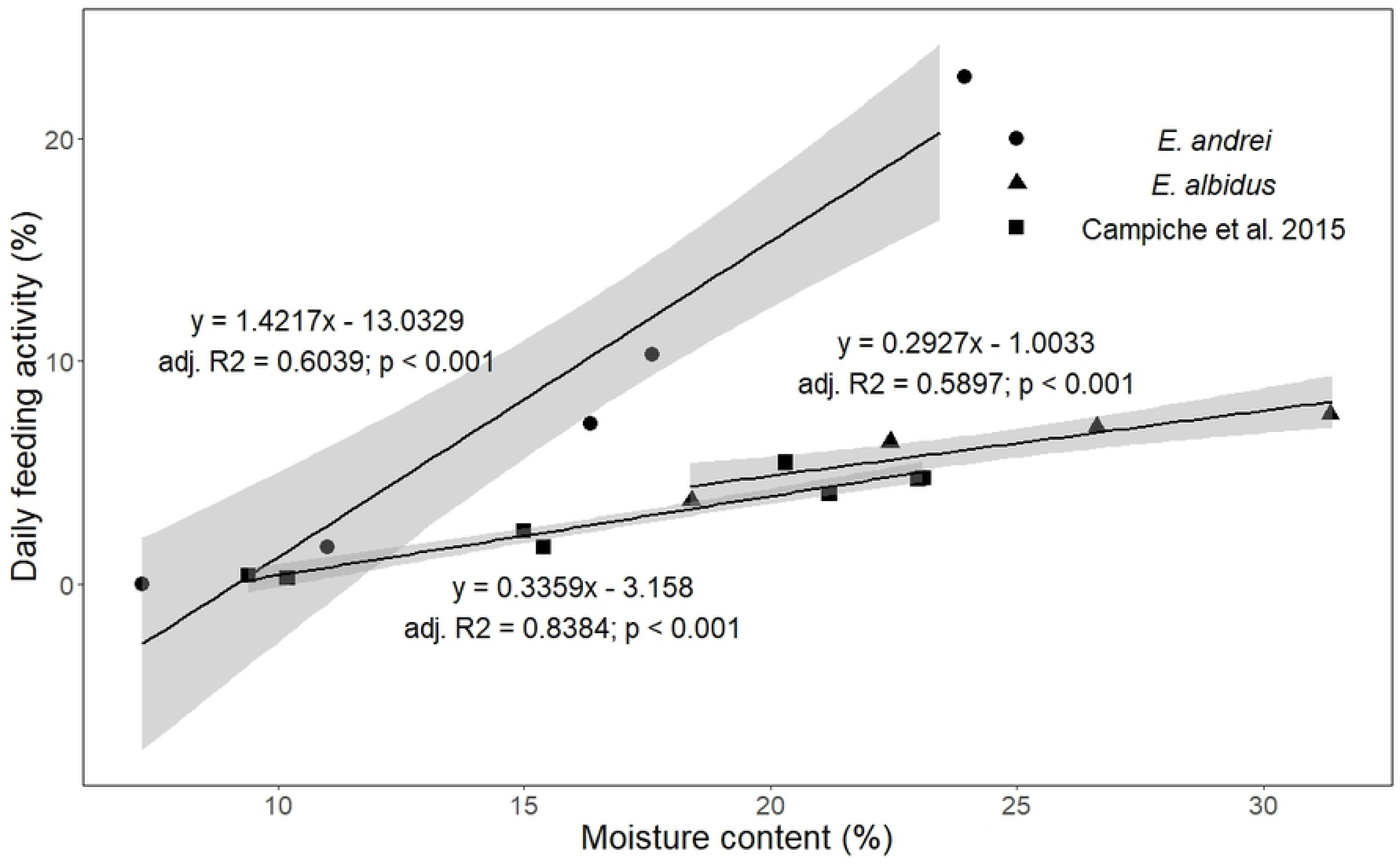
Linear regressions (black lines) and 95% confidence interval (grey shades) of the daily feeding activity (in % of pierced holes) of each experiment exposed to different soil moistures (gravimetric water content, %). Each symbol (black dots for *E. andrei*, this study, black triangle for *E. albidus*, this study, black squares for Campiche et al. 2015 [20] represents the average daily feeding activity per treatment.

**Table 4.** Parameters of the multiple regression analysis for the daily feeding activity response to the two predictors ’moisture’ and ’experiments’, with interaction between the two factors. Data on moisture are expressed as gravimetric moisture content (%). The experiments compared are *Eisenia andrei* (this study), *Enchytraeus albidus* (this study), and Campiche et al. [20]. β0, β1, β2, and β3 are the coefficients of the linear regression. Adjusted R square = 0.6051.

| 1) $y = \beta_0 + \beta_1 \cdot \text{moisture} + \beta_2 \cdot \text{experiment} + \beta_3 \cdot \text{moisture} \cdot \text{experiment}$ | | | | |
| --- | --- | --- | --- | --- |
| *E. andrei* VS Campiche et al. [20].<br>Adjusted R<sup>2</sup> = 0.6759 |  |  | *E. albidus* VS Campiche et al. [20].<br>Adjusted R<sup>2</sup> = 0.8489 |  |
| Coefficient | Estimate (95% confidence interval) | p value | Estimate (95% confidence interval) | p value |
| $\beta_0$ | -13.0329 (-18.2403 – -7.8254) | < 0.001 | -1.003 (-3.6296 – 1.6231) | 0.525 |
| $\beta_1$ | 1.4217 (1.1252 – 1.7181) | < 0.001 | 0.2927 (0.1865 – 0.3990) | < 0.001 |
| $\beta_2$ | 9.8474 (1.9266 – 17.7682) | < 0.05 | -2.1822 (-5.0632 – 0.6989) | 0.134 |
| $\beta_3$ | -1.0657 (-1.5111 – -0.6203) | < 0.001 | 0.0620 (-0.0618 – 0.184) | 0.313 |
| *E. albidus* VS Campiche et al. [20].<br>Adjusted R<sup>2</sup> = 0.8486 |  |  |  |  |
|  | Estimate (95% confidence interval) | p value |  |  |
|  | -2.1118 (-3.5594 – -0.6643) | < 0.01 |  |  |
|  | 0.3384 (0.2824 – 0.3944) | < 0.001 |  |  |
|  | -0.7709 (-1.4906 – -0.0511) | < 0.05 |  |  |

## 4. Discussion

### 4.1. Test performance and optimal moisture conditions for the two species

Test replicates were in general drier at the end of the tests compared to the start, and the differences were more important for the earthworm test. The lower variation in moisture between test start and test end found for enchytraeids compared to earthworms can be explained by the fact that, for this species, the humidity was monitored and adjusted during the experiment, because of the longer test duration. A humidity check during the test, as suggested in standard guidelines for testing reproduction, is thus recommended for the bait lamina test in the laboratory.

The bait lamina test is rarely performed under extreme soil moisture conditions. Recently, their suitability under saturated media has been showed by means of sediment assays [16] and the study demonstrates the promising potential of using bait lamina to assess organic matter breakdown in the sediment compartment. In the present study, no bait loss was recorded in negative control replicates without test organisms with the only exceptions of the two highest tested moistures (nominal 80 % and 100 % WHC), where a slight bait loss was observed (see SI2). Even though this was probably due to bait dissolution by water, the bait loss in both cases was always very low (maximum 2 % over the whole test duration), suggesting that globally the bait lamina test can be used under a very wide range of soil moisture contents, from very dry to more than saturated.

In general, both *E. andrei* and *E. albidus* were confirmed to be efficient feeders of bait material, with an average daily feeding activity at the reference moisture content of 22 % and 6 %. Because a similar biomass per replicate was used in the two experiments, when comparing the results at the two reference moisture contents, it can be concluded that *E. andrei* was more than three time faster than *E. albidus* in consuming the bait material.

For earthworms, feeding activity was highest (i.e., 21.82 ± 10.41) at the reference moisture content (i.e., 52 % WHC), suggesting that for bait lamina test, the optimal moisture conditions are in the same range recommended for reproduction guidelines of *E. fetida/andrei* (40 to 60 % WHC) [22,23] and close to the recommended value (60 % WHC) in the guideline for assessing avoidance behavior [21]. Daily feeding activity measured at the reference soil moisture was comparable to values found in other laboratory experiments, performed with same or similar species (*Eisenia andrei/fetida*) and at similar soil moisture contents. Van Gestel et al. [7] exposed ten earthworms in approximately 814 g of OECD artificial soil with 5 bait lamina strips and estimated that the time required to empty 50 % of the holes corresponded to approximately 4.6 days. Casabé et al. [14] obtained an average feeding activity of 62.8 % after three days (i.e., 22.7 % of consumed bait per day) by exposing six earthworms in 350-400 g of uncontaminated clay silty soil with 4 bait lamina sticks. Finally, Jänsch et al. [15] obtained more than 30 % feeding activity after seven days when exposing 10 earthworms in artificial and eight natural soils with 2 bait strips. When normalized to test duration, number of earthworms, and number of bait sticks used, the results of feeding activity correspond to 0.95 [14]; 0.46 (this study); 0.22 [7]; and 0.21 [15] % of consumed bait per day per earthworm per bait stick. Mesocosms or terrestrial model ecosystems with natural soil [4–6] suggested as well a high contribution of earthworms to bait consumption, but overall feeding activity was in general lower compared to values observed in the above mentioned laboratory experiments. This difference can be explained by the fact that earthworm’s species other than *E. andrei/fetida* were present in the experiments but also by the complexity resulting from test conditions, which were closer to a field situation. Particularly, the use of intact soil cores soil structure is maintained, contrarily to classical single-species laboratory experiments, where sieving of the soil leads to structure loss. Other reasons can also be different intrinsic properties of the tested soil, such as organic matter content, texture, presence of other food sources that could have been preferred over the bait material, or interaction within soil communities.

For enchytraeids, the optimal soil moisture for feeding activity was at 68 % WHC (7.62 ± 0.39 % of consumed bait per day), so at a higher moisture content than the reference (49 % WHC). However, the results were not significantly different for these two moisture treatments, suggesting that the range recommended in the standard guidelines (between 40 and 60 % of maximum WHC [24,25]) can be suitable for use in bait lamina tests. The slightly higher optimum for feeding activity for enchytraeids compared to earthworms might suggest a higher tolerance for greater levels of moisture contents. Like for earthworms, our experiments confirmed that enchytraeids are important consumers of the bait, which is aligned with findings of other laboratory studies. Helling et al. [29] exposed 75 adults enchytraeids (*E. minutus* and *E. lacteus*) in 75 g of arable soil and artificial OECD soil for 10 days with one bait lamina stick and obtained 17 and 7 % of feeding activity, respectively. Bart et al. [13] observed approximately 70 % of feeding activity after 5 days of exposure of *E. albidus* in 50 g of a luvisol with 1 stick. When normalized to test duration, number of enchytraeids, and number of bait sticks used, the results of feeding activity correspond to 1.16 [13]; 0.23 [29]; and 0.01 (this study) % of consumed bait per day per earthworm per bait stick. The higher results of feeding observed in the literature compared to this study could be partially explained by the fact that a lower amount of soil was used in the mentioned studies compared to this study: if less soil is used, there could be a higher probability of access to the bait-lamina as well as a reduction of alternative food sources. Differently, a laboratory experiment, showed very low feeding activity of the enchytraeid *Cognettia sphagnetorum* which was exposed in a forest soil, under a wide range of soil moistures and temperatures [4]. Such lower activity compared to other laboratory experiments might be explained by the very high content in soil organic matter, which could have been a preferred source of food for those enchytraeids rather than the bait. A similar hypothesis was formulated by André et al. [30] who recorded a significant negative correlation between soil organic matter content and feeding activity for a site characterized by a thick litter layer.

### 4.1 Moisture effect on feeding activity

For both test species, feeding activity was low with weak soil moisture contents and increased with growing soil water contents until a peak feeding activity was reached at the species-specific optimum moisture level. In general, it is well acknowledged that moist soils are more favorable for earthworm activity [31] and that, similarly, enchytraeids are sensitive to drought [32–34].

Unlike low moisture, the effect of very high moisture contents was different for the two species. For earthworms, at moisture levels higher than the optimum (i.e., 52% WHC), the average feeding activity decreased but remained relatively constant and without a clear correlation to soil water content. Despite high moisture conditions negatively affecting the performance of earthworms [35], many species are able to survive even for long periods in soil totally submerged by water [31]. Similar observations could be found in our study, where *E. andrei* survived at all relative moisture contents up to 104 % WHC but with reduced feeding activity relative to their optimum.

For enchytraeids, there was only one moisture treatment higher than the optimum (i.e., 69% WHC) and thus, there are not enough data available to investigate possible correlations between feeding activity and high soil moisture. At this highest moisture treatment, enchytraeids feeding activity was not only significantly reduced compared to the reference, but was almost absent. In this treatment, the soil had a flooded appearance and many enchytraeids, while active and alive, remained on the soil surface. The behavior observed for enchytraeids suggests that they were less able to burrow into flooded soil compared to earthworms and it was then more difficult for them to reach the bait strips. Despite having a higher optimum, *E. albidus* seems thus to be more strongly impacted than *E. andrei* by moisture contents above the optimum.

The increase in feeding activity with increasing soil moisture could be modelled by linear regressions for soil moisture contents but only up to the optimum. However, the considered range is expected to cover optimal moisture conditions for monitoring feeding activity, which are suggested to be between slightly moist to field capacity [36]. For soils having a similar texture to LUFA 2.2 soil (i.e. sandy loam), plant wilting occurs at a gravimetric water content lower than 7 %, while field capacity is situated around 20 % [37]. The models derived for earthworms and enchytraeids cover a range between 7 to 23 % and 18 to 31 % gravimetric water content, respectively (see SI2). It could thus be suggested that both models would be relevant for optimal monitoring conditions of soil moisture contents, for sandy loamy soils at expected field capacity. However, care should be taken when comparing field soils and sieved soils because of possible changes due to structure loss and further testing is needed to verify this hypothesis.

The multiple regression performed for the two tested species showed a significant interaction between species and soil moisture content, meaning that the increase in feeding activity over the defined soil moisture range was significantly different between the two species. This increase was more pronounced for *E. andrei* compared to *E. albidus*, which is not surprising, as the first has been shown to feed on the bait faster than the latter. Different factors could explain the faster bait consumption of *E. andrei* compared to *E. albidus*. One explanation is purely biological, and assumes that earthworms simply have an inherent higher feeding rate than enchytraeids. A second hypothesis is mechanical and could be that earthworms, being bigger than enchytraeids, have more facility to break and feed on the bait material. Indeed, enchytraeids and earthworms play similar functional roles, but at different spatial scales [38]. Finally, there might be other reasons related to the resources and intrinsic habitat quality of the LUFA 2.2 soil, which might have been more attractive for *E. albidus* than for *E. andrei*. A limited amount of resources in the soil for *E. andrei* could make that they would feed preferentially on the bait material. On the other hand, *E. albidus* could have found sufficient resources in the LUFA 2.2 soil during the first days, which would explain why they fed more slowly on the bait material. This last hypothesis could be supported with the fact that earthworms and enchytraeids have been suggested to have slightly different ecological preferences and niches [39,40].

The comparison made between this study and the previous field experiment suggests that the model developed for *E. albidus* (this study) is more similar, compared to *E. andrei*, to the model developed in the field study Campiche et al. [20]. This could suggest that in the arable field investigated by Campiche et al. [20], the bait lamina consumption is driven more by the enchytraeid community rather than by earthworms. It is worth mentioning that the species used in this study (*E. andrei*) is not a relevant representative of agricultural fields [38]. In addition to species, earthworms can have different feeding behaviors depending on which ecological group they belong. Epigeic species (to which *E. andrei* belongs) live and feed more on litter material and are considered to initiate the decomposition process of complex organic material. On the other hand, endogeic species prefer pre-processed organic material, while anecic earthworms mainly transport organic litter from the soil surface to deeper soil layers [41]. It is not known if and how each of these ecological groups contribute differently to bait consumption. From our results, it can be suggested that epigeic earthworm species are not the best representative of the agricultural field studied by Campiche et al. [20], whereas *E. albidus* seems to better reflect the biological activity of the field. This is in line with some observations suggesting that in agricultural ecosystems *Enchytraeus* species are generally more abundant [42] and more ecologically relevant than earthworms, such as *E. andrei* [38,43]. It cannot be excluded that other species and/or ecological categories than *E. andrei* and/or epigeic species were present and active in the field from Campiche et al. [20], and that they may have had a smaller contribution to bait consumption. In general, for each site studied, further investigations would be needed to assess the real soil community and draw more consistent conclusions on the relative contribution of each specific soil organism group to feeding activity, measured through the bait lamina test.

Globally, the results of our study are an important first step for normalizing bait-lamina test data between studies and sites based on soil moisture, which is one of the biggest confounding factors in field testing. So far, *E. albidus* seems to be a more pertinent indicator for this kind of comparison, because of the similarity in the slopes with the field condition study. However, to further validate the produced model, additional factors should still be explored, which can have an influence on feeding activity under field conditions. Intrinsic soil properties, such as texture or organic matter content, can strongly influence water retention and therefore the activity of soil organisms [29,44]. The bait lamina test should be performed with other soil types than the LUFA 2.2. to explore potential differences in the model. In addition, the soil used in laboratory experiments is generally sieved, disrupting the original structure that can be found under natural field conditions. The loss of structure can have an important influence on the capacity of the soil to retain water and therefore typical characteristic such as wilting point and field capacity might be less comparable. Environmental factors (e.g., temperature) are also known to play a role on the behavior of the soil community as well as soil conditions. Finally, natural communities of soil organisms are more diverse and complex than the limited number of species that can be tested in the laboratory and this can influence the velocity of bait consumption as well as the main feeders of the bait, as already highlighted by the two tested species.

## Conclusions

The bait-lamina method has been shown to be a suitable tool for evaluating the impact of soil moisture on feeding activity of two soil species used in standard testing. For both earthworms and enchytraeids, feeding activity increased linearly with increasing soil moisture up to an optimum. In addition to species differences in feeding rates and optimum moistures, the linear increases in feeding activity were also found to be species dependent. The linear model was able to describe the feeding response of two important soil invertebrate species for a realistic range of soil moisture contents and *E. albidus* seemed to be provide more similar outcomes compared to field data. The produced models will facilitate interpretation of future field studies, but first additional studies are required to validate this model under complex field conditions.

## Acknowledgements

We would like to thank INRAE/AgroParisTech (UMR ECOSYS, Versailles, France) for having kindly provided the species *Enchytraeus albidus* Henle 1847. We would also like to thank the Statistical Consulting service, at the ETH of Zurich, for their help in the statistical analysis.

## Supporting information captions

**S1 Table. Field abundances of earthworms for arable soils in Switzerland and neighboring countries**

**S2 Table. Field abundances of enchytraeids for arable soils in Switzerland and neighboring countries**

**S3 Table. Soil moisture content, feeding activity and mortality data for bait-lamina tests on earthworms and enchytraeids**

**S4 Figure. Comparison of nominal and measured soil moisture contents for bait-lamina tests with earthworms in the first (A) and second (B) test**

**S5 Figure. Comparison of nominal and measured soil moisture contents for bait-lamina tests with enchytraeids**

